# Modeling human embryo adhesion using a microfluidic platform

**DOI:** 10.64898/2026.03.10.710513

**Authors:** Sofía Zaragozano, María Pardo-Figuerez, Ana Monteagudo-Sanchez, Javier Gonzalez-Fernandez, Javier Moncayo-Arlandi, Anna Quirant, Sara Maggi, Luis Quintero, Francisco Raga, Juan Pablo Grases, Xavier Santamaria, Inmaculada Moreno, Nicolas Plachta, Carlos Simon, Felipe Vilella

## Abstract

Embryo adhesion represents the first step of implantation, yet understanding this process has been hindered by the lack of human in vitro platforms that replicate endometrial physiology. Here, we present a dual-channel microfluidic platform containing organoid-derived endometrial epithelium and primary stromal cells. Our model recapitulates important endometrial hallmarks including epithelial polarization, stromal decidualization, extracellular vesicle release, and hormone-induced receptivity. We tested its function by introducing mouse and human blastocysts and showed that embryos displayed features of initial adhesion. These included establishment of embryo-epithelial contacts initiated via the polar trophectoderm, inner cell mass repositioning, and lineage segregation. Moreover, human embryos secreted βhCG indicating a functional trophoblast. Thus, this work provides a platform to study key features of embryo adhesion and endometrial receptivity, and disorders affecting embryo-endometrium interactions.

**Teaser:** Endometrium-on-a-chip shows detailed human embryo adhesion dynamics.

## Introduction

Embryo implantation is the essential first step for establishing a pregnancy and involves blastocyst apposition, adhesion to the endometrium, and invasion into the decidua (*1*). Adhesion serves as gatekeeper of the periconceptional period influencing long-term health outcomes for both mother and child (*1*). Insights into human embryo adhesion derive primarily from in vivo studies compiled into the Carnegie stages of human development, a foundational reference in human embryology (*2*). Complementary studies in murine models have elucidated dynamic changes in cell–cell adhesion and cytoskeletal architecture that enable the morphogenetic remodeling of the trophectoderm, critical for the successful transition of the blastocyst into the post-implantation embryo (*3–5*). However, significant physiological differences between mouse and human implantation may affect the direct applicability of these findings, especially regarding the mechanism of embryo adhesion (*6*).

Initial in vitro models based on human endometrial epithelial cells (EECs) (*7*) or immortalized EEC lines (*8*) have shifted to 3-dimensional (3D) systems (*9*). The development of endometrial epithelial organoids (EEOs) has recently offered the ability to replicate 3D architecture and hormone response (10, 11). Although insightful, these models lack endometrial stromal cells (ESCs), essential for the crosstalk between the embryo and endometrium in the in vivo environment (*10–13*). To address this, 3D systems have incorporated EEOs within 3D matrices that support co-culture with stromal cells and other cell types (*14–16*). Yet, these systems lack dynamic flow and have architectural complexity which difficult embryo implantation studies.

Microfluidic devices offer control over cellular interactions, chemical gradients and mechanical forces, which are crucial for mimicking in vivo physiology (*17*, *18*). These systems have proven useful for modeling hormone response (*19*), microbiome-host interactions (20, 21), and implantation dynamics within the female reproductive tract (*22*). Building on these advances, we developed an endometrium-on-a-chip model using a commercially available microfluidic platform to investigate adhesion using both mouse and human embryos. Our design replicates endometrial architecture and cellular communication allowing the study of embryo adhesion.

## Results

### Development and characterization of an in vitro adhesion dynamics on a chip model (ADOC)

To investigate the mechanisms underlying embryo adhesion in vitro, we first developed an endometrium-on-a-chip model by using a commercial microfluidic device consisting of two channels connected by a 7 µm-pore membrane, allowing communication between upper and lower compartments. We named this model ADOC (Adhesion Dynamics On a Chip). ADOC design is based on an epithelial compartment in the upper channel of the system and a stromal compartment in the lower channel (Fig. 1A and B). To establish the epithelial compartment, organoid-derived EECs were grown as a monolayer in the top channel while stromal cells were seeded from primary cultures (see Material and Methods). Following six days of culture, both channels showed confluency (Fig. 1B and fig. S1). Immunofluorescence analysis confirmed that EECs expressed keratins, a marker of epithelial identity (*23*) (Fig. 1C), while ESCs exhibited the expression of the mesenchymal marker vimentin (Fig. 1C). F-actin staining revealed the epithelial monolayer distribution and highlighted the characteristic spindle-shaped morphology of stromal cells (fig. S1). Visualization of the chip organization in 3D showed that epithelial cells arranged in a columnar structure, similar to their in vivo counterparts (*24*) (Fig. 1D). Furthermore, stromal cell migration through the porous membrane towards the epithelial compartment suggested interaction across the channels (Fig. 1D).

**Fig. 1.**
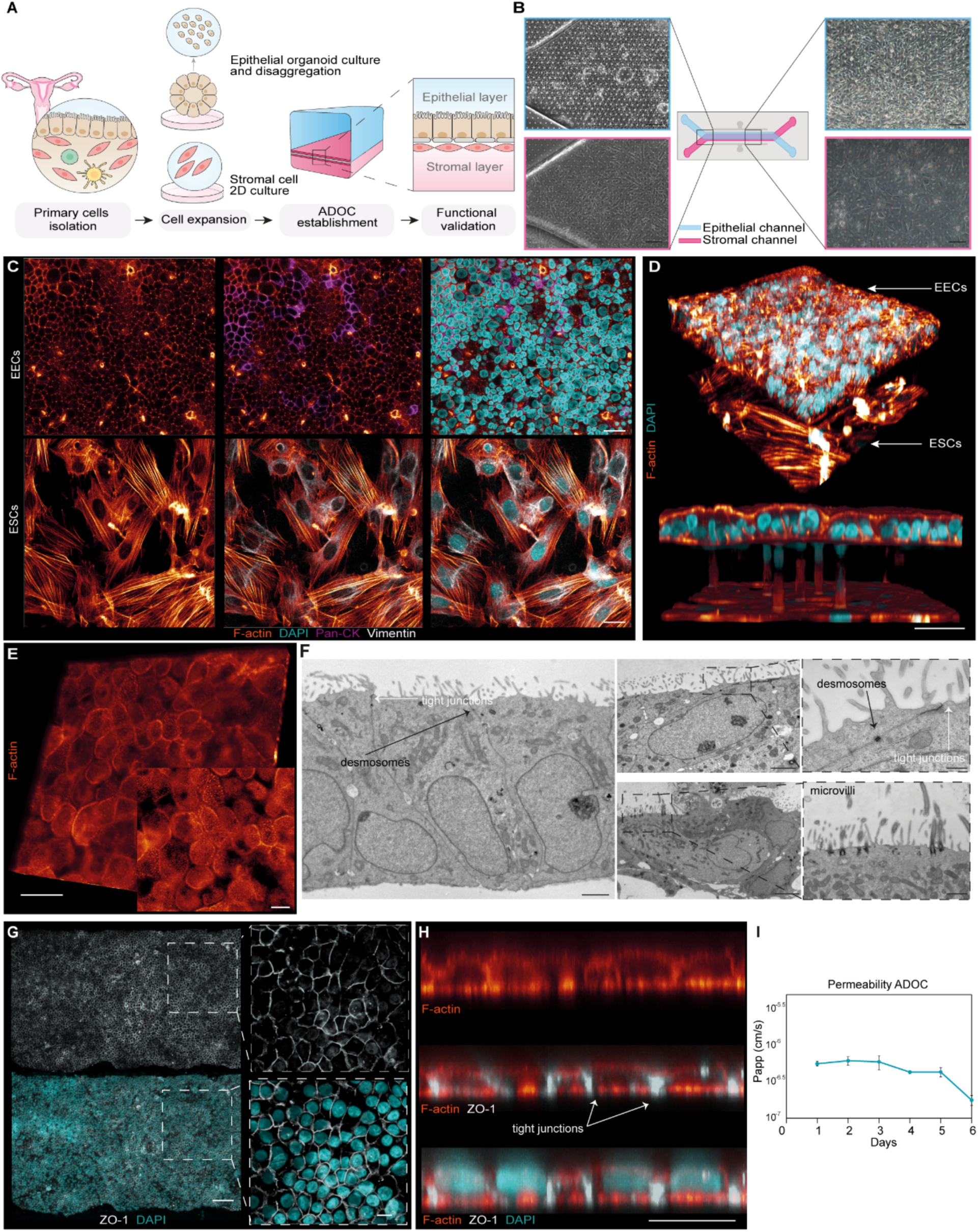
Development and characterization of an in vitro Adhesion Dynamics On a Chip model (ADOC). **(A)** Schematic overview of the establishment of the endometrium-on-a-chip model. **(B)** Diagram and brightfield images showing the epithelial (blue) and stromal (pink) channels after six days of growth. Scale bars, 100 µm. **(C)** Immunostaining of epithelial cells (top) with F-actin (orange), DAPI (blue), and Pan-Cytokeratin (purple) and stromal cells (bottom) with F-actin (orange), DAPI (blue), and Vimentin (white). Scale bar, 20 µm. **(D)** Top: 3D projection showing the epithelial monolayer (above) and the stromal layer (below) stained with F-actin (orange) and DAPI (blue). Bottom: A blend rendering showing the transversal cross-section of ADOC highlights the columnar morphology of the epithelium and migration of stromal cells toward epithelial cells. Scale bars, 40 µm. **(E)** Epithelial compartment stained with F-actin revealed microvilli as distinct dots. Scale bars, 100 µm. **(F)** Electron microscopy shows desmosomes (black arrows) and tight junctions (white arrows), confirming epithelial cell polarization. Scale bars, 2 µm, 500 nm. **(G)** Immunostaining of tight junctions (ZO1-gray) demonstrates a well-formed epithelial barrier. Scale bar, 100 µm. **(H)** Cross-section of the epithelial layer highlights the tight junction (ZO1-gray) deposition within individual cells. Scale bars, 100 µm. **(I)** Permeability assay using cascade blue shows minimal molecular exchange after cell growth in the model (data is presented as the mean ± S.D.).

Characterization of the epithelial compartment showed apical microvilli in the EEC monolayer, visualized by F-actin (Fig. 1E). Transmission electron microscopy (TEM) confirmed epithelial cell polarization, revealing basally located nuclei surrounded by organelles, and an apical surface densely covered with microvilli (Fig. 1F). We further identified tight junctions and desmosomes (Fig. 1F), crucial for epithelial polarity (*25*). Immunofluorescence for the tight junction protein ZO-1 revealed that cell membranes were in tight contact forming a continuous barrier structure (Fig. 1, G and H), essential for preserving epithelial integrity (*26*). To evaluate the stability and integrity of the epithelial barrier, we measured the apparent permeability (P_app_) over 6 days with cascade blue, a standard fluorescent tracer for paracellular permeability in microfluidic models (*27*). P_app_ values remained stable between day 1-4, ranging from approximately 1.2 × 10⁻⁶ to 8.0 × 10⁻⁷ cm/s, indicating early formation of a functional barrier. By day 6, P_app_ decreased markedly, reflecting an enhanced barrier maturation over time (*28*, *29*) (Fig. 1I).

### Hormonal response to epithelial receptivity and stromal decidualization

To investigate ADOC functionality, we assessed hormone response to epithelial receptivity and decidualization. After 6 days of culture, both channels of ADOC were treated with 17β-estradiol (E2) for 2 days, followed by a combination of E2, medroxyprogesterone acetate (MPA), cyclic adenosine monophosphate (cAMP) and the Tankyrase Inhibitor XAV-939 for an additional 4 days (Fig. 2A & Material and Methods). These components facilitate stromal decidualization while inhibiting the WNT/β-Catenin pathway to create a receptive epithelium (*13*).

**Fig. 2.**
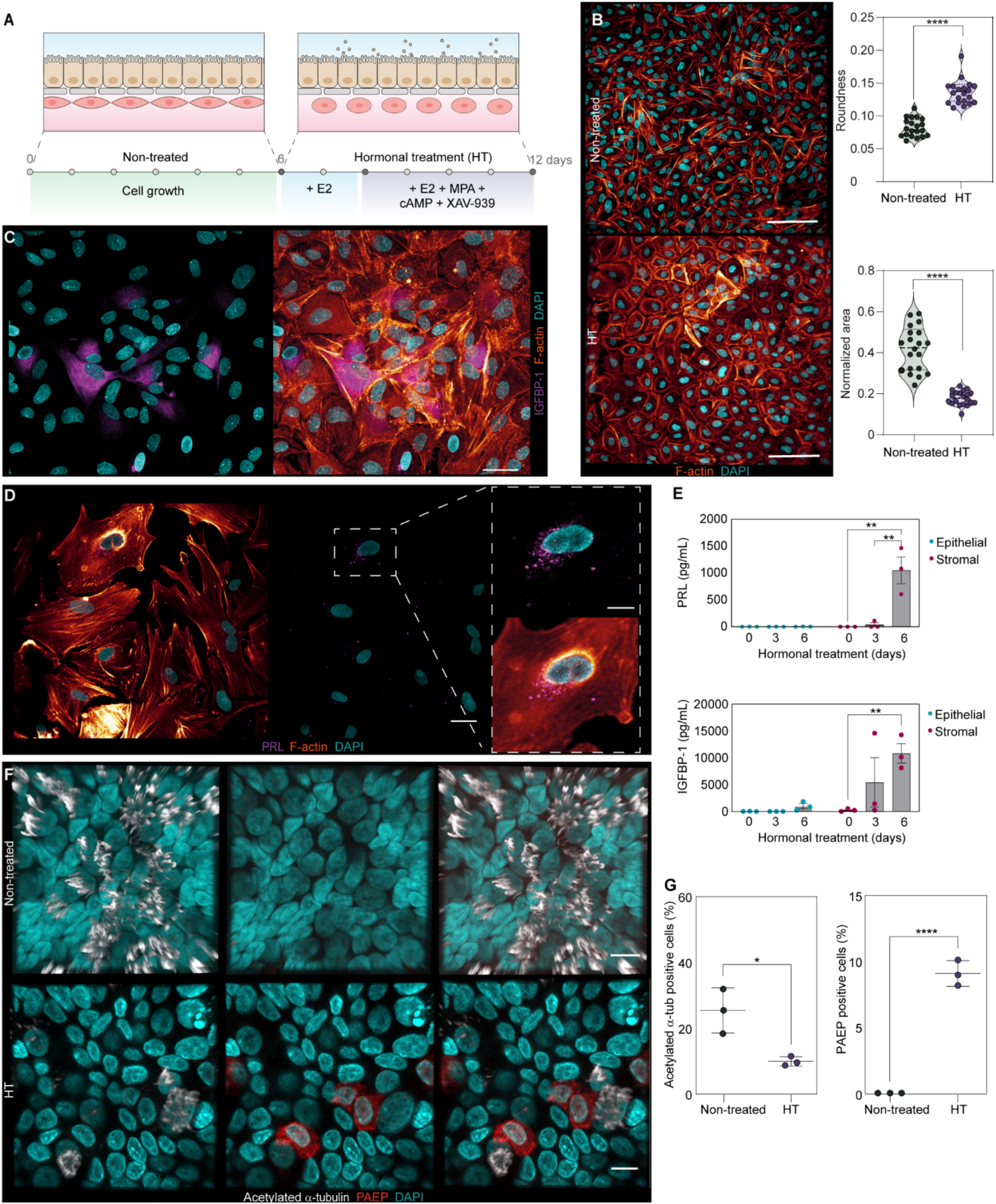
Epithelial receptivity and stromal decidualization in ADOC. **(A)** Schematic of the experimental setup. **(B)** Decidualization of stromal cells after six days of hormonal treatment. Left: Immunostaining of stromal cells before and after hormonal treatment with F-actin (orange) and DAPI (blue). Right: Morphological quantitative analysis showing increased roundness (top panel) and decreased cellular area (bottom panel) of stromal cells after hormonal treatment. Data is presented with violin plots, ****p<0.0001 by Mann-Whitney test and unpaired t-test respectively. Scale bars, 70 µm. **(C, D)** Presence of IGFBP-1 (purple) and PRL (pink) in response to hormonal treatment, along with DAPI (blue) and F-actin (orange) staining. Scale bars, 50 µm. **(E)** PRL and IGFBP-1 quantification using ELISA at days 0, 3, and 6 of hormonal treatment. Data is presented as mean ± S.D., n=3, **p<0.01 by Bonferroni’s multiple comparisons 2-way ANOVA. **(F)** Acetylated α-tubulin (white) and PAEP (red) in endometrial epithelial cells before and after hormonal treatment. Scale bar, 10 µm. **(G)** Quantification of the percentage positive ciliated and PAEP cells before and after hormonal treatment. Data is presented as mean ± S.D., n=3, *p<0.05, ****p<0.0001 by unpaired t-test.

In the stromal compartment, after six days of treatment, cells transitioned from spindle to rounded shape (Fig. 2B), in agreement with the literature. Quantitative analysis revealed a reduction in cell area and increase in roundness, compared to non-treated cells, consistent with stromal decidualization remodeling (*30*) (Fig. 2B). Hormone treatment also induced the secretion of IGFBP-1 and PRL, accompanied by PGR expression, suggesting a functional hormone response and decidualization (Fig. 2, C and D and fig. S2A). Enzyme-linked immunosorbent assay (ELISA) assay for IGFBP-1 and PRL performed with the culture supernatant during hormonal treatment showed no detectable levels on day 0, an increase by day 3, and a further rise by day 6 (Fig. 2E).

The upper epithelial monolayer also responded to hormonal stimulation, as evidenced by a reduction in acetylated α-tubulin-positive ciliated cells (25.4 ± 3.9% vs. 9.4 ± 0.8%). This response was accompanied by an increase in PAEP/glycodelin-positive cells following treatment (0% vs. 9.5 ± 0.5%) (Fig. 2, F and G). The hormonal treatment also affected the permeability barrier, as evidenced by an increase in P_app_, which is consistent with the permeability changes previously described (*31*, *32*) (fig. S2B). These findings suggest that the upper epithelial channel becomes receptive and that ADOC replicates aspects of the secretory endometrium.

### ADOC recapitulates endometrial extracellular vesicle secretion

We next examined whether the model recapitulates the endometrial extracellular vesicle (EV) secretion, a process known to play a key role in maternofetal crosstalk (*33–35*). TEM micrographs revealed multivesicular bodies in epithelial (Fig. 3A) and stromal cells (Fig. 3B). To confirm EV secretion, we collected media from both compartments between days 5 and 11, which revealed the three major EV subtypes including apoptotic bodies (ABs), microvesicles (MVs), and exosomes (EXOs) (Fig. 3, C and E and fig. S3A). To determine differences in the size and concentration of MVs and EXOs derived from both compartments, we performed nanoparticle tracking analysis (fig. S3, B, and C). Our analysis revealed significant differences between epithelial- and stromal-derived MVs and EXOs. Specifically, the epithelial layer secreted larger EVs in greater quantities (Fig. 3, D and F and fig. S3, B and C). Comparison of epithelial and stromal EVs before and after hormonal treatment showed no differences suggesting that EV release is maintained independent of hormonal treatment (fig. S3D), in line with previous work (*35*). To test that the in secretory phase EVs resembled those in vivo, we extracted their miRNA content and analyzed the presence of endometrial-specific miRNAs including hsa-let-7e-5p and hsa-miR-17-5p typically enclosed within EVs (*36*). qRT-PCR analysis confirmed that we were able to detect them at similar cycle thresholds to endogenous miRNA expressed in EVs (Fig. 3G), supporting that ADOC recapitulates the EV profile of the endometrium.

**Fig. 3.**
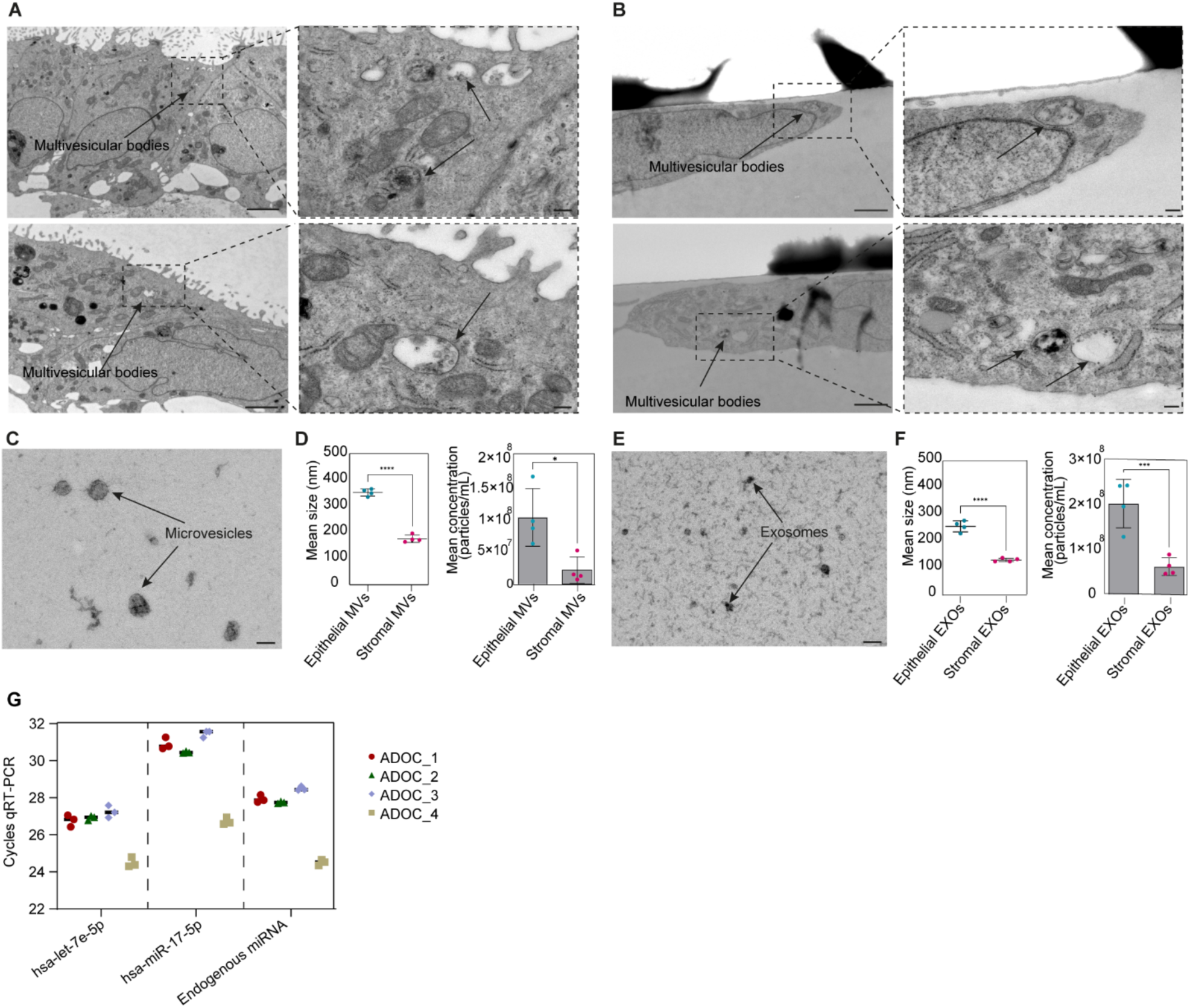
ADOC recapitulates endometrial extracellular vesicle secretion. **(A)** Transmission electron microscopy (TEM) micrographs of epithelial cells showing multivesicular bodies localized in the apical region of the cells. Scale bar, 2 µm (left), 200 nm (right). **(B)** TEM micrographs of stromal cells displaying multivesicular bodies. Note that the empty spaces and black structures correspond to the membrane of ADOC. Scale bar, 1 µm (left), 200 nm (right). **(C)** TEM micrograph of microvesicles (MVs) isolated from ADOC. Scale bar, 100 nm. **(D)** Nanoparticle tracking analysis (NTA) quantification of mean size (left) and concentration (right) of epithelial vs. stromal MVs isolated from the epithelial and stromal compartments after six days of hormonal treatment (data is presented as mean ± S.D., n=4, ****p<0.0001, *p<0.05, by unpaired t-test). **(E)** TEM micrograph of exosomes (EXOs, arrows) isolated from ADOC. Scale bar, 100 nm. **(F)** Nanoparticle tracking analysis (NTA) quantification of mean size (left) and concentration (right) of epithelial vs. stromal EXOs isolated from the epithelial and stromal compartments after six days of hormonal treatment (data is presented as mean ± S.D., n=4, ****p<0.0001, ***p<0.005, by unpaired t-test). Scale bar, 500 nm. **(G)** RT-qPCR detection of selected microRNAs (let-7e-5p, miR-17-5p, and endogenous control) in epithelial-derived MVs isolated from four independent replicates (ADOC_1–4).

### Mouse embryos display adhesion in ADOC

We then explored whether ADOC could be used to study embryo adhesion. Mouse blastocysts were introduced into the epithelial channel following hormonal treatment (Fig. 4A). A total of 23 mouse blastocysts (52%, n=44) successfully adhered to the epithelial compartment, confirmed by their resistance to media flushing (Movie S1). Mouse embryo attachment progressed through an initial adhesion stage (“adhesion onset”), characterized by the firm attachment of the first embryonic cells to the epithelial layer, followed by the spreading of the embryo across the epithelium (Fig. 4A). The average time at which adhesion occurred was 59.3 ± 29.5 hours (fig. S4A). During initial adhesion, the cells forming the embryo appeared densely packed while maintaining its spherical shape (Fig. 4B). Following this, the embryo gradually spread out over the epithelial surface, increasing contact area (“adhesion spreading”) (Fig. 4B). To quantify this morphological change, we measured embryo spreading area over the epithelial monolayer, revealing an increase in adhesion area compared to the onset of this process (Fig. 4C). Immunofluorescence for the trophectoderm (TE) marker GATA3 confirmed that trophectoderm cells initiated contact, while the inner cell mass (ICM) labeled by OCT3/4 remained apically positioned (Fig. 4D, fig. S4B and Movie S2). In the spreading adhesion stage, we could confirm that TE cells spread and flattened over the epithelial layer (Fig. 4E). The analysis of trophectoderm spreading using computational analysis confirmed that the majority of trophectoderm nuclei (GATA3) aligned within the same focal plane of the epithelium layer (DAPI EECs) along the Z-axis, suggesting a full spreading across the epithelial layer (Fig. 4, F and G). Furthermore, a blastocoel cavity restructuring was observed (Fig. 4H), suggesting ICM polarization. After full spreading, we also observed hypoblast reorganization, as revealed by the presence of GATA4 localized closer to the EECs than OCT3/4 cells (Fig. 4, I and J).

**Fig. 4.**
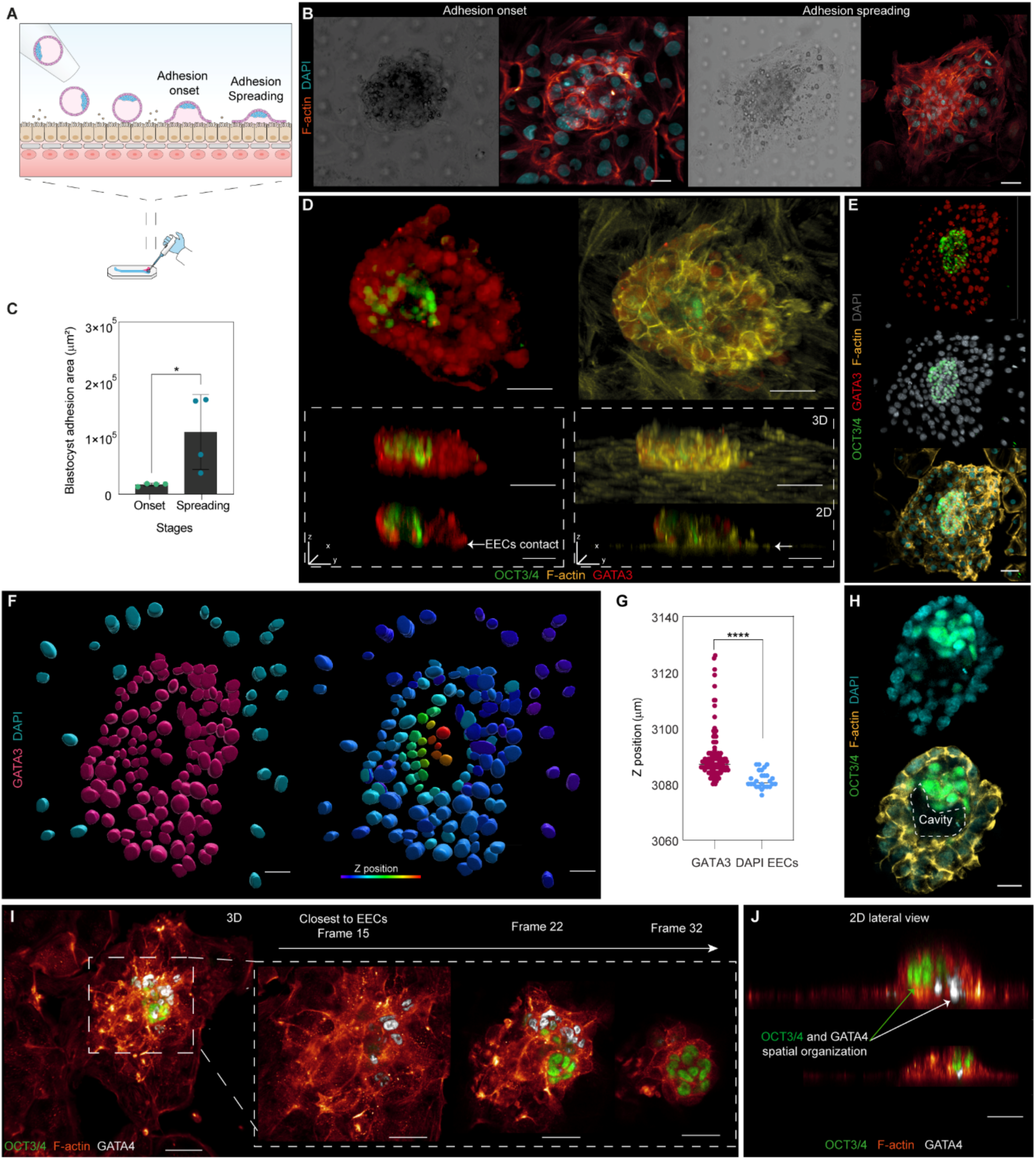
Mouse embryo adhesion confirms functionality of the ADOC model. **(A)** Schematic of embryo positioning in ADOC showing embryo placement on the epithelial layer and the progression from adhesion onset to spreading. **(B)** Representative images of mouse blastocysts during adhesion onset and spreading. Brightfield images (left) and immunofluorescence staining for F-actin (orange) and DAPI (blue) reveal changes in cell morphology and cytoskeletal organization. Scale bars, 40 µm. **(C)** Quantification of blastocyst adhesion during adhesion onset (blastocyst collapse) and spreading, demonstrating significant increase during adhesion progression (Data are shown as mean ± S.D.; unpaired t-test, *p < 0.05). **(D)** Representative immunofluorescence images (top panel) of adhered blastocysts highlighting the inner cell mass (ICM; OCT3/4, green) and trophectoderm (GATA3, red), along with F-actin (yellow). Middle panel: 3D reconstruction of blastocyst adhesion in the epithelial compartment. Bottom panel: 2D Y-Z planes showing the spatial positioning of the inner cell mass (ICM; OCT3/4, green) and trophectoderm (GATA3, red) TE. White arrows indicate the onset trophectoderm contact with the epithelial layer. Scale bars, 50 µm (top), 80 µm (bottom, 3D), 40 µm (bottom, 2D). **(E)** Fluorescent images of a fully expanded blastocyst and surrounding epithelial cells. OCT3/4 (green), GATA3 (red), F-actin (yellow), and DAPI (blue) suggest full spreading across the layer. Scale bar, 50 µm. **(F)** Computational segmentation of fully spread embryo showed the GATA3 nuclei position in a close plane as epithelial layers (DAPI EECs). Color scale indicates Z position: blue = closer to epithelium, red = farther from epithelium. Scale bar, 30 µm. **(G)** Dot plot showing Z position distribution of GATA3 nuclei and epithelial DAPI cells. **(H)** Representative immunofluorescence images of adhered blastocysts (top panel), highlighting the inner cell mass (ICM; OCT3/4, green), DAPI (blue), and F-actin (yellow). Dotted lines indicate the presence of a cavity. Scale bar, 20 µm. **(I, J)** Mouse blastocyst showing lineage specification after adhesion in ADOC. **(I)** 3D reconstruction and sequential optical sections (left, Frame 15 closest to epithelial cells) to upper planes (right, frame 22 and 32) reveal that GATA4 endoderm cells (white) are located more adjacent to the epithelial cell layer, whereas OCT3/4 epiblast cells (green) are positioned apically, slightly away from the epithelium. Scale bars, 50 µm (3D), 40 µm (sections). **(J)** 2D lateral view showing epiblast and primitive endoderm positioning. OCT3/4 epiblast cells (green) are located apically, while GATA4 primitive endoderm cells (white) are situated closer to the epithelial interface Scale bar, 50 µm.

### ADOC reveals adhesion dynamics and early lineage reorganization of human blastocysts

We finally studied adhesion of human blastocysts (day 5-6 post-fertilization (p.f.), Fig. 5A). Following the selection of hatched human blastocysts, we used time-lapse brightfield imaging to record the dynamic process of embryo positioning and adhesion (Fig. 5, A and B). During the adhesion process, the embryo underwent a morphological transition from a spherical, expanded blastocyst to a flattened compacted structure. Pre-adhesion images showed a well-defined blastocoel and rounded morphology across both planes, whereas post-adhesion images revealed a loss of sphericity, increased optical density, and close contact with the epithelial surface in the lower plane (Fig. 5B, Movie S3). Out of 19 blastocysts, 47% (n=9) attached over a mean period of 34.1±12.2 hours (Fig. 5C).

**Fig. 5.**
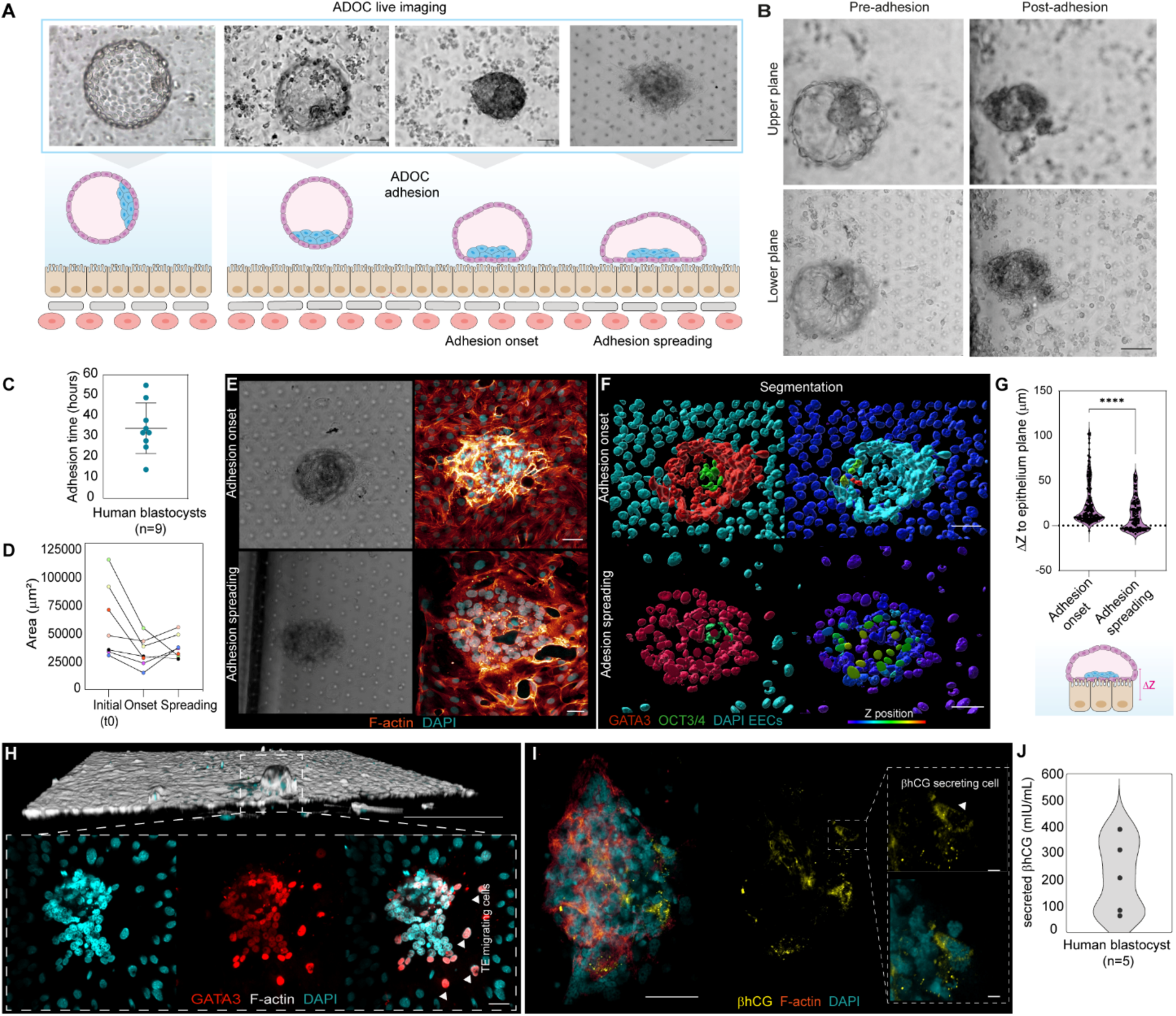
Human blastocyst adhesion and invasion dynamics in the ADOC model. **(A)** Time-lapse sequence of a human blastocyst adhering to the epithelial layer in ADOC. Top: brightfield time-lapse images show the morphological changes during different stages of adhesion. Bottom: schematic representation of each stage seen in vitro. **(B)** Time lapse images showing upper (focusing the embryo) and lower (focusing the epithelial layer) planes before (pre) and after (post) adhesion into the layer. Scale bar, 50 µm. **(C)** Average time of adhesion onset for individual blastocysts. Data are shown as mean ± SD (n=9). Each dot represents a different human embryo. **(D)** Quantification of embryo area in ADOC at three stages: initial placement (t₀), onset of adhesion, and spreading. Each line represents an individual embryo, and colored points are connected by lines to indicate temporal area evolution for each embryo (n=7). **(E)** Representative immunofluorescence images of adhesion onset and spreading stages. Left panels: brightfield images. Right panels: F-actin staining (orange) highlights cytoskeletal organization. DAPI (blue) marks nuclei. Scale bar, 50 µm. **(F)** 3D computational segmentation of embryos at adhesion onset and spreading stages. GATA3 (red) marks trophectoderm cells, OCT3/4 (green) marks the ICM and DAPI (blue) marks epithelium. **(G)** Quantification of embryo position relative to the epithelial layer during adhesion. Violin plots represent the Z-distance (ΔZ, in µm) between the embryo and the epithelial reference plane during two distinct adhesion phases: Adhesion onset and spreading. ΔZ was calculated as the distance from the centroid of the embryo surface to the average Z-plane of the epithelial nuclei. Positive values indicate displacement above the epithelium. (Dots indicate individual nuclei, ****p < 0.0001, Mann–Whitney U test). Bottom schematic illustrates ΔZ quantification relative to epithelial monolayer. **(H)** 3D side view of an adhered embryo during the spreading stage in ADOC. Zoom-in reveals trophectoderm (GATA3, red) extending along the epithelial surface. Arrowheads indicate individual trophectoderm cells migrating laterally. Scale bars, 3D:100 µm, 50 µm. **(I)** Representative immunofluorescence images demonstrating βhGC release after implantation. Scale bars, 50 µm, 10 µm (zoom). **(J)** βhCG secretion by individual human blastocysts (n=5). Violin plot represents the distribution of secreted βhCG (mIU/mL), with dots indicating individual embryo values.

Quantitative tracking of embryo area over time showed a decrease coinciding with blastocyst collapse indicative of adhesion onset (Fig. 5D). This was followed by an increase in embryo area in most of the cases, reflecting trophoblast spreading across the epithelial layer, as also observed with F-actin staining (Fig. 5, D and E). 3D reconstructions of segmented embryos allowed to identify Z position of each nuclei and showed that at the onset of adhesion, most embryo cells (GATA3) were positioned above the epithelial nuclei, as identified by DAPI staining (and GATA3-cells). By contrast, during spreading, trophectoderm shifted closer to the epithelial nuclei, indicating increased integration with the epithelial surface (Fig. 5F). Quantitative analysis confirmed that the onset of adhesion displayed larger ΔZ values whereas the adhesion spreading exhibited lower ΔZ values (Fig. 5G). Additionally, late spreading showed lateral migration of TE cells towards the epithelial layer. Although ADOC is designed to study embryo adhesion, this early migratory event could suggest the onset of embryo invasion (Fig. 5H).

To study functional trophectoderm activity, we demonstrated that adhered embryos released β-human chorionic gonadotropin (βhCG) confirming that adhesion was coupled with βhCG hormone secretion by trophoblast cells (Fig. 5I), recapitulating early events of embryo adhesion (*37*). Measured βhCG concentrations in the culture media after adhesion confirmed the presence of the hormone in the media which was variable for each embryo (Fig. 5J).

We further explored spatial cell arrangement during human blastocyst adhesion. Optical sections from different planes revealed that the lower plane, adjacent to the epithelium, was primarily composed of GATA3 TE cells, but also included a few internally positioned OCT3/4 cells, all merged across the epithelial layer (Fig. 6A, below). In the upper plane, OCT3/4 cells were compactly clustered and surrounded by trophectoderm cells (Fig. 6A, above), suggesting that the ICM remained largely internal but still in close contact with the epithelial layer during the first steps of adhesion (Fig. 6A and Movie S4). Higher magnification into the ICM (Fig. 6B) confirmed a radially organized OCT3/4 cell cluster surrounded by GATA3 cells, suggesting the first steps of lumen formation, indicative of embryo initial reorganization (*38*).

**Fig. 6.**
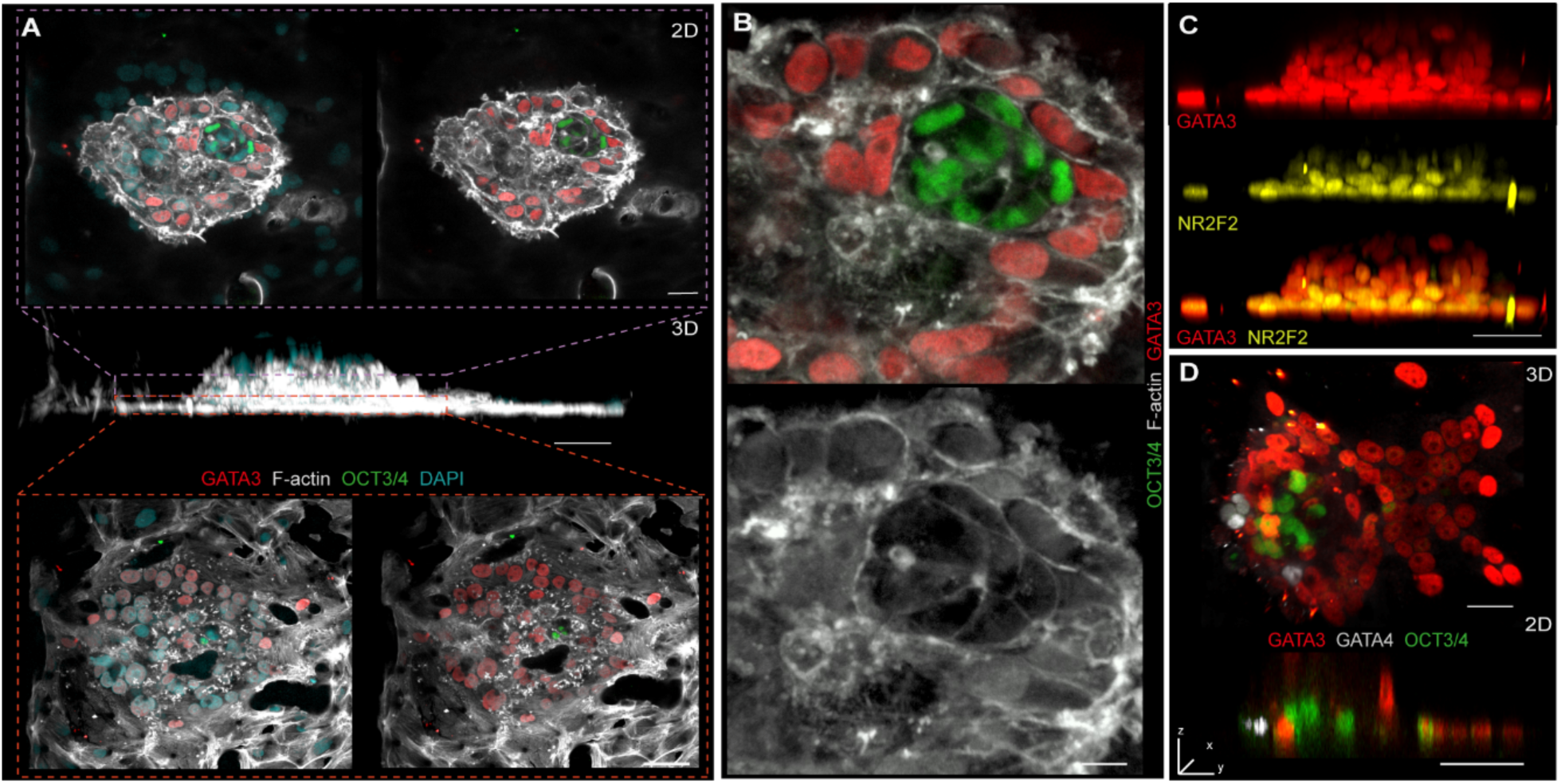
Human blastocyst adhesion in ADOC reveals trophectoderm spatial localization and ICM remodeling. **(A)** 3D projection of a blastocyst during adhesion. Zoom-in of optical sections at two different focal planes— one closer to the epithelial layer (below, orange dashed lines) and one farther away (above, purple dashed lines) demonstrating that few OCT3/4 cells positioned adjacent to the epithelial surface, while GATA3 TE cells occupy from basal to apical positions. Scale bars, 50 µm(3D), 30 µm. **(B)** The cluster of OCT3/4 cells (green) representing the ICM are centrally located and surrounded by GATA3 cells (red). The OCT3/4 cells form a compact, radially organized structure, as shown with F-actin (white) Scale bar, 15 µm. **(C)** 3D side views of adhered blastocysts stained for OCT3/4 (green), GATA3 (red), and NR2F2 (yellow). Scale bars, 50 µm. **(D)** 3D projection showing three distinct embryonic lineages: GATA4 primitive endoderm (white), OCT3/4 epiblast (green), and GATA3 trophectoderm (red). Scale bars, 30 µm, 50 µm.

To confirm that embryos adhered via the polar trophectoderm, we examined the localization of NR2F2, a marker of polar TE identity in the human blastocyst. NR2F2 cells were localized at the region of the TE in contact with the epithelial surface, confirming that embryo adhesion in ADOC occurs via the polar TE (Fig. 6C), thus supporting correct embryo orientation for adhesion and developmental progression. To study further lineage specification, staining for GATA4 revealed additional ICM complexity, with GATA4 primitive endoderm cells reorganizing with OCT3/4 epiblast and GATA3 TE populations (Fig. 6D). These three lineages were segregated in 2D orthogonal views demonstrating lineage specification during early human implantation stages. These sections showed that a layer of GATA4 and OCT3/4 cells were consistently positioned closer to the epithelial layer along with GATA3 cells, which surrounded the periphery and extended apically (Fig. 6D). Altogether, our model effectively captures the dynamics and spatial organization of early human embryo adhesion and enables controlled study of embryonic–maternal interactions.

## Discussion

The early stages of human implantation, including adhesion, remain poorly understood due to ethical constraints and the inherent challenges of accessing these events in vivo. Current assays include in vitro human endometrial models, which range from 3D primary tissue constructs to organoid-based and microfluidic systems. Some of these models support co-culture with either human embryos or blastoids up to the adhesion stage (*13–15*, *39*), but the key features used to establish embryo adhesion still remain difficult to study. To address this, we developed ADOC, in which hormonal treatment successfully mimicked the mid-secretory phase endometrium, inducing stromal decidualization and epithelial receptivity (*40–43*). Although hormone response has been previously described in vitro (*10*, *16*), our model relies on independently cultured epithelial and stromal cells, each perfused with dynamically flowing medium. Although advantageous, this design display some limitations since the porous membrane acts as a physical barrier to direct embryonic invasion into the stromal compartment. However, the modular design of the chip provides flexibility to incorporate additional cellular compartments. Indeed, endothelial cells have been successfully integrated into a three-channel endometrium-on-a-chip model (*22*); however, this model was developed using cell lines and was not tested for embryo functionality.

With the current design, we demonstrated critical aspects of human embryo adhesion. We successfully captured the processes of human blastocyst apposition and adhesion, visualizing distinct morphological transitions such as polar trophectoderm spreading and ICM repositioning under hormonally-receptive conditions. Previously reported 3D models (*14*, *15*) have provided valuable insights into human implantation. However, limited imaging depth and architectural complexity made it difficult to resolve embryo–epithelium interactions with sufficient spatial resolution. Whereas initiation of adhesion by the polar trophectoderm has been reported in vitro (*44*), our model uniquely enables visualization of this process alongside the self-organization of the ICM into a radial pattern, comparable to early lumenogenesis observed in human embryos (*44*, *45*). Furthermore, we demonstrated reorganization of hypoblast-like cells during adhesion spreading. While these observations align with previous in vitro studies of embryo differentiation and implantation assays using blastoids (*13*, *15*), our work provides the first evidence of both lineage specification and the initiation of lumenogenesis in a human preimplantation blastocyst within an endometrial context.

In conclusion, our microfluidic endometrium-on-a-chip platform enables real-time imaging and precise tracking of human embryo adhesion. Future studies could use this platform for functional genomics, host–microbiome interaction studies, and pharmacological screening to identify compounds that could improve embryo adhesion impairment. Given that endometrial diseases and implantation failure remains a major challenge in assisted reproduction (*46*, *47*), ADOC offers a powerful tool to investigate the mechanisms underlying these pathologies.

## Materials and Methods

### Experimental design

A commercial microfluidic device (Emulate) with two interconnected channels was employed to develop an endometrium-on-a-chip model for studying embryo adhesion. Endometrial biopsies were collected from patients under ethical approval, and primary cells were processed to separate epithelial and stromal fractions. The epithelial cells were cultured as organoids, while stromal cells were maintained in a 2D culture system. The organoids were dissociated and seeded into the upper channel of the microfluidic device, while stromal cells were seeded into the lower channel. Both cell types were expanded under continuous flow for six days, with organoid and stromal medium, respectively. To mimic the secretory phase of the endometrium, the model was treated with E2 for two days, followed by a combination of E2, cAMP MPA and XAV939 for four days.

To test the functionality of the model, mouse blastocysts (n=44) were devitrified, assisted-hatched, and cultured overnight in G2 plus® medium. Two hours before implantation, the blastocysts were transferred to EmbryoGlue® and placed in the epithelial channel of the device, where adhesion was monitored using confocal microscopy with time-lapse imaging. To study human adhesion dynamics, human blastocysts (n=19; day 5–6 post-fertilization) donated for research were devitrified, assisted-hatched, and cultured for two hours in embryo medium before being transferred to EmbryoGlue® and cultured overnight. The human embryos were then introduced into the epithelial channel and monitored for adhesion events using confocal microscopy.

Mouse and human embryos were fixed at different stages of adhesion and characterized using immunofluorescence to evaluate molecular markers and adhesion progression. This experimental design allowed the assessment of the model’s ability to support and study embryo adhesion under conditions that simulate the human endometrium.

### Samples and ethical approvals

All tissue samples utilized in this study were obtained with written informed consent from all participants, in full compliance with the ethical guidelines set forth in the Declaration of Helsinki 2000. Superficial endometrial biopsies were collected as part of the IGX1-ORG-FV-21-01 project, which was approved by the Clinical Research Ethics Committees of Clínica El Pilar (Barcelona) and Hospital Clínico Universitario de Valencia. Eligibility criteria for participation in the study included female patients aged 18–42 years, within the reproductive age range, with a natural menstrual cycle and a body mass index (BMI) between 18.5 and 29.9 kg/m². Participants with oncological conditions, intrauterine device (IUD) use within the past three months, hormonal treatment within the previous month, or bacterial or viral infections were excluded from the study. Human embryos donated for research as surplus from IVF treatments were obtained from Next Fertility clinics. Their use was approved by the Ethics Committee on Drug Research (CEIM) of the Hospital Universitario y Politécnico La Fe (Valencia), as well as by the National Commission on Assisted Human Reproduction and the Commission on Guarantees for the Donation and Use of Human Tissues and Cells (Spain), under the framework of project FCS-IMA-CS-22-2.

### Processing of endometrial biopsies

A two-stage dissociation protocol was used to desegregate endometrial biopsies into stromal fibroblast and epithelial-enriched single-cell suspensions (*43*). Firstly, the sample was rinsed with phosphate-buffered saline (PBS) (Biowest, L0615) in a petri dish to remove blood and mucus. Then, the tissue was minced into small pieces and dissociated with collagenase V (Sigma, C9263), RPMI Gibco, 21875-034), 10% fetalbovine serum (FBS, Biowest, S181B-500) and DNAsa I (Sigma, 11284932001) at 37°C for 25 min under continuous shaking (175 rpm). The content was transfer to falcon through a 100 μm cells strainer filter. The tissue remaining on the filter was used for epithelial enrichment by incubating with 10 mL trypsin-DNAse I for 10 min in the previous conditions. The resulting two contents were transferred to 50 mL with 20 mL RPMI and filtered with 100 μm cell strainers.

### Endometrial organoids

Endometrial organoids were generated using previous described protocol (*10*) with minor modifications. The epithelial fraction obtained from the biopsies was resuspended in 30% DMEM/F12 (Thermo Fisher Scientific, 11330032) and 70% Matrigel (Corning, 356231) and supplemented with the rho-associated protein kinase, Rock inhibitor (RI, Y-27632, Merck SCM075), at a final concentration of 2µM. The suspension was cultured in 20 μL droplet deposited in prewarmed 48-well plates. Organoids were cultured as previously described (see Table S1) (*48*). Matrigel droplets were cultured for 15 days in the first passage (P0), and for 7 days in the following passages. The medium was changed every 2 days. Organoids were recovered for passaging by liquifying Matrigel droplets with ice-cold DMEM/F12. Then, the organoids were dissociated using TrypLE Select (Gibco, 12563-029) for 7 min at 37°C supplemented with 1 μL RI and mechanically triturated. The resulting cells suspension was centrifugated at 300 x g and resuspended in 70% Matrigel and 30% DMEM.

### Endometrial stromal cells culture

The stromal fraction was seeded in T-25 flasks using DMEM-F12-Glutamax (Gibco, 10565) media supplemented with 10% FBS, 0.2% Gentamycin (Gibco, 15750037) and 0.2% Amphotericin B (Cultek, 5530-003-CF) for expansion and selection of fibroblastic-like cells. The cells used were from early passages (P0 to P5).

### Human pre-implantation embryos culture

Human donated embryos (day 5-6 post fertilization) were thawed using Cryotop safety thawing kit following the manufacturer’s instructions (Kitazato, VT602) and underwent a 30% of assisted hatching (Octax eyewere laser, Vitrolife). Embryos were cultured in 30 μL droplet of Embryo medium(*15*) (Table S2) covered with Hypure oil light (Kitazato, 96005) for 2 hours, then embryos were cultured in EmbryoGlue® (Vitrolife, 10085) overnight at 37°C and 5% CO_2_.

### Mouse embryos culture

Mouse blastocysts (Embryotools®, PRO-005) were thawed following manufacturer’s instructions (Cryotop safety thawing kit, Kitazato, VT602) and underwent a 30% of assisted hatching. Blastocysts were cultured in 30 µL of G2 plus® (Vitrolife, 10132) droplets cover with Hypure oil light overnight at 37°C and 5% CO_2_. Then, blastocysts were transferred to EmbryoGlue® for 2 hours before its placement in the upper channel of the chip.

### Establishment of primary human endometrium-on-a-chip

The ADOC model is based on a commercially available platform (Emulate), in which we culture EEOs cells and primary stromal endometrial cells. To functionalize the chip, ER-1 (Emulate, 10461) and ER-2 (Emulate, 10462) were mixed to reach a 1 mg/mL concentration and added to the top and bottom microfluidic channels, subsequently the chip was irradiated with UV light having a peak wavelength of 365nm (NailStar^TM^, NS-01-EU) for 20 minutes. Then, both channels were coated with tissue-specific ECM components, 100 µg/mL collagen type I (Merck, C3867-1VL), and 25 µg/mL fibronectin (Merck, 11080938001) for stromal cells and 100 µg/mL Matrigel (Corning, 356231) for epithelial cells. The coated chip was incubated 2 hours in the incubator at 37°C followed by a 4°C overnight incubation. The next day, EEOs were disaggregated, and single epithelial cells were seeded (1×10^5^ cells/ 35µL confluence) on the top channel. The following day, ESCs were seeded (1×10^5^ cells/ 15µL confluence) on the bottom channel. Chips were connected to the Zoë® instrument and perfused continuously at a flow rate of 30 µL/h, using expansion media (EM) consisting in organoid and stromal media, respectively.

### Hormonal treatment of endometrium-on-chip

After 6 days of growing in expansion medium, the epithelial cells derived for EEOs and the stromal cells seeded on the chip, were treated for 2 days with 10 nM E2 (Sigma-Aldrich, E2758) to mimic the proliferative phase, and followed by the mixture of 10nM E2, 1µM MPA (Sigma, M1629), 250 µM cAMP (Sigma, B7880) and 10 µM XAV939 (Deltaclon, S1180) for 4 days to mimic the secretory phase.

### In vitro implantation assay

After the six days of hormonal treatment, ADOC epithelium was prepared at least 2h prior the placement of blastocysts by washing the channel two times with 200 μL of EmbryoGlue® and kept at 37°C. The embryos were transferred to the upper channel using a 200 μL pipette under an inverted microscope. The chips were placed over a Ibidi plate (Ibidi, 82107) to perform live imaging. To confirm embryo adhesion, PBS was flushed into the upper channel of the chip. Embryos that remained adhered following the flushing were used for subsequent analyses.

### Immunofluorescence staining

Firstly, upper and down channels were washed with 200 μL of PBS three times, and then cells were fixed with 4% paraformaldehyde (FisherScientific, 28908) for 30 min at room temperature. After fixing, the channel was washed three times with PBS. The cells were permeabilized in PBS 0.5% or 1% Tween-20 (Sigma, P9616) between 1-3 hours, and block with PBS 0.1% Tween-20 5% BSA (Myltenic biotec, 130-09-376) for 1 hour, all the steps were performed at room temperature in agitation. The samples were then incubated overnight at 4°C in agitation with primary antibodies (Table S3) diluted in blocking solution. The day after, the samples were washed with PBS 0.1% Tween-20 at least three times. The washing buffer was replaced by the secondary antibodies (Invitrogen) (Table S4I) diluted in blocking buffer and incubated between 2-3 h at room temperature in agitation. Then, the samples were washed at least three times with PBS 0.1% Tween-20. To finish a last incubation with DAPI (Merck, D9542) and/or phalloidin (ab235137, ab176759, Abcam; A34055, Invitrogen) diluted in PBS-BSA 1% was performed between 30 min-1h. Finally, the channel was washed with PBS for three times.

### Transmission electron microscopy

For electron microscopy studies, samples were fixed with 3% glutaraldehyde in 0.1 M phosphate buffer, and membranes were extracted from the chip. Samples were post-fixed with 2% osmium, rinsed, dehydrated, and embedded in Durcupan resin (Fluka, Sigma-Aldrich, St. Louis, USA). Ultrathin sections (0.06-0.08 µm) were made with an Ultracut UC-6 (Leica microsystems, Wetzlar, Germany) with a diamond knife, stained with lead citrate (Reynold’s solution), and examined under a transmission electron microscope FEI Tecnai G2 Spirit BioTwin (ThermoFisher Scientific Company, Oregon, USA). All images were acquired with a Xarosa digital camera (EMSIS GmbH, Münster, Germany) controlled by Radius software (Version 2.1).

### Enzyme-linked immunosorbent assay IGFBP-1 and PRL

Conditioned media from the top and bottom outlets were collected on days 0, 3, and 6 of hormonal treatment. PRL (Boster biological technology, EK0593) and IGFBP-1 (Raybiotech, ELH-IGFBP1-1) concentrations were assayed using commercial ELISA kits according to the manufacturer’s instructions. A 1:20 dilution of the media was used for testing IGFBP-1 and a dilution 1:2 of the media was used for tested PRL. The IGFBP-1 test limits detection to 5 pg/mL and the PRL test to 10 pg/mL. Each test was quantified by measuring optical density at 450 nm in a Multiscan Go (ThermoScientific).

### ELISA assay for βhCG detection

The media from epithelial channel was collected at various time points following human embryo adhesion and stored at -20°C until use. The medium was tested for βhCG levels using βhCG ELISA (Abcam, ab1786339) according to manufacturer’s instructions, alongside with βhCG standards. The results were quantified by measuring optical density at 450 nm in Multiscan Go (ThermoScientific).

### Permeability assay

To evaluate the permeability of the epithelial barrier formed in the chip we added Dextran, Cascade Blue™, 3000 MW (ThermoScientific) (*49*) to the top channel inlet reservoir at 100 µg/mL diluted in expansion organoid media using a flow rate of 30 µl/h. Culture media from the top and bottom inlet reservoirs and top and bottom outlet reservoirs were sampled every 24 h after chip connection to the Zoë® perfusion system. The fluorescence intensity of Cascade Blue™ was measured at 400/420 nm excitation/emission wavelengths using a Victor III plate reader (PerkinElmer, Inc, Waltham, USA).

The apparent paracellular permeability (P_app_) was calculated based on a standard curve using the following formula:

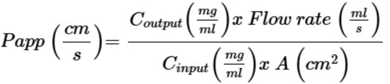

Where: C_output_ is the concentration of Cascade Blue™ in the effluents of the stromal compartment, A is the seeded area of epithelial channel, and C_input_ is the input concentration of dextran spiked into the epithelial compartment

### Extracellular Vesicle Isolation

EVs were isolated from conditioned media collected from top and bottom outlet reservoirs in non-treated and hormonal treated chips. Stromal culture media supplemented with 10% FBS contains a variable amount of small extracellular vesicles; therefore, EVs were removed from FBS by ultracentrifugation at 120,000 x g for 70 min at 4°C using a P50AT4 rotor in a CP100NX centrifuge (Hitachi) to ensure the isolation of only EVs from stromal cells within the chip. The supernatant containing EV-free-FBS was used to supplement the stromal media for this experiment. Isolation of EVs from effluent culture media was performed as previously described with minimal modifications (*34*). Culture medium was centrifuged at 300 x g for 10 min to pellet residual cells and debris. Supernatants were centrifuged at 2,000 x g for 10 min to separate and pellet the apoptotic body (AB) fraction. Pellets were resuspended in 1 mL of cold PBS and centrifuged at 2,000 x g for 10 min to obtain a clean AB fraction. Supernatants were passed through a 0.8 µm diameter filter (Whatman plc Maidstone) and centrifuged at 10,000 x g for 40 min to pellet the fraction enriched in MVs. Pellets were resuspended in 1 mL of cold PBS and centrifuged at 10,000 x g for 40 min to obtain a clean MV fraction. Finally, supernatants were passed through 0.22 µm diameter filters (Acrodisc® syringe filters, Pall Corporation) and centrifuged at 120,000 x g for 70 min to pellet the EXO fraction of the media. Pellets were resuspended in 1 mL PBS and centrifuged at 120,000 x g for 70 min to obtain a clean exosome-enriched fraction. All centrifugation steps were conducted at 4°C, and pellets for ABs, MVs, and EXOs after the PBS centrifugation step were resuspended in 50 µL PBS and saved at -80°C.

### Nanoparticle tracking analysis of isolated EVs

EV size distribution and concentration in the effluent media of chips were measured by nanoparticle tracking analysis (NTA) principles in a NanoSight 300 instrument (Malvern Instruments Corp.). Due to the limited size working range (<1,000 nm particle), ABs could not be analyzed. 10 µL of the resuspended pellet of EVs were diluted in 700 µL PBS and introduced into the NanoSight 300. The starting media volume for EV isolation was used to normalize concentration measurements.

### miRNA qPCR analysis

miRNA isolation from MVs obtained from collected media was performed using the miRNeasy micro kit (Qiagen, 217084) following the manufacturer’s instructions. For the isolation the ath-miR159a (5’phos-UUUGGAUUGAAGGGAGCUCUA-3’; IDT) was spike-in as exogenous control. Total isolated RNA was quantified using the QuickDrop instrument (Molecular Devices). Complementary DNA (cDNA) synthesis of miRNAs was performed using TaqMan® Advanced miRNA cDNA Synthesis Kit (ThermoFisher Scientific, A28007) from 2 µL of total isolated RNA following the manufacturer’s instructions. Real-time qPCR of miRNAs was performed using TaqMan^TM^ Fast Advanced Master Mix (ThermoFisher Scientific, 4444557) and specific probes for each miRNA (ThermoFisher Scientific) in a total volume reaction of 10 µL in a 384-well-plate and a QuantStudio 5 Real-Time PCR System (ThermoFisher Scientific).

The expression of hsa-let-7e-5p (A25576, 478579_mir), and hsa-miR-17-5p (A25576, 478447_mir) was evaluated. The miRNAs, has-miR-200c-3p (A25576, 478351_mir) and hsa-miR-92a-3 (A25576, 477827_mir) were used as endogenous controls. To normalize data, we used exogenous controls. To evaluate the expression of miRNAs in the MVs, we compared the cycle threshold (Ct) values of the target miRNAs to those of ubiquitously expressed endogenous controls.

### Microscopy and image analysis

The phase-contrast, fluorescent images and time-lapse images were acquired using Leica Stellaris 5 and software associated (LASX Version 4.8.0.28989). The 3D reconstructions and 2D planes were analyzed using FIJI 1.53k or Bitplane Imaris 9.7.0 software. For live imaging, embryos were scanned every hour at the upper, lower and middle plane of the embryo on the chip using a 10x objective.

Confocal immunofluorescence images of mouse and human blastocysts were acquired with a 10x CS×20/0.75 numerical aperture air-immersion or a 25x objective numerical aperture water-immersion objective. For immunofluorescence images, optical sections from 0.57 to 1.5 µm were acquired.

In Figure 2B, morphological changes of ESCs before and after decidualization were analyzed by changes in area and roundness. For this, images taken at random areas of the stromal compartment from three different chips were taken at 25x. For each cell, the area and roundness were measured manually outlining the cell boundary in ImageJ. Roundness was calculated by ImageJ as indicated in:

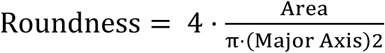

with values approaching 1 indicating more circular shapes.

In Figure 2G, positive cells for PAEP and acetylated α-tubulin were identified based on specific marker expression and divided by the total number of DAPI-stained nuclei within each field of view. Data is presented as the percentage of marker-positive cells relative to the total cell number. Blastocyst adhesion time was determined by continuous monitoring of human embryos cultured on the microfluidic chip. Embryos were imaged at regular intervals to identify the timepoint at which adhesion onset occurred, defined as the moment of blastocoel collapse and stable contact with the epithelial layer. To validate adhesion, embryos were fixed in situ at different timepoints. Adhesion was confirmed when embryos remained immobile upon application of the fixative, indicating sufficient attachment to resist flow-induced displacement.

Blastocyst adhesion area in mouse and human was quantified by manually outlining the perimeter of each embryo in 2D slicers. The outer boundary of each embryo was manually delineated at the plane providing the best visibility of the contact region with the epithelial layer, and the corresponding surface area was calculated using the software’s measurement tools. For human blastocyst, “initial” indicated the area of the blastocyst once introduced in the chip. For mouse and human blastocyst, “adhesion onset” was defined morphologically during live imaging as the stage at which the blastocoel collapsed and optical density changed, indicating initial attachment. We referred to “adhesion spreading” when blastocyst expansion and lateral spreading across the epithelial surface of ADOC was observed.

To evaluate spatial positioning of embryo and epithelial cells, the tool surface was used to manually segment the nuclei in Imaris. Epithelial nuclei were segmented from DAPI cells lacking GATA3 expression, while embryonic nuclei were identified as GATA3/DAPI. The Z position (depth) of each segmented nucleus was extracted from the centroid coordinates provided by Imaris. These values were used to compare the vertical distribution of embryonic (GATA3) and epithelial nuclei and assess whether they occupied similar focal planes during embryo adhesion.

For human embryos, quantification of ΔZ distance to epithelium was measured by assessing the relative position of the embryo within the microfluidic device. For this, we generated 3D surfaces of the epithelial nuclei (DAPI) and the embryo (GATA3) and analyzed the ΔZ distance as the difference between the Z position (Z size of 0.57 µm) of the embryo and that of the average nuclei position in epithelial monolayer. Positive ΔZ values indicate that the embryo was located above the epithelial layer, while values close to zero indicate contact or integration with the epithelium.

## Statistical analysis

Statistical analyses were performed using Graphpad prism 8.1.1. Normality of the data distribution was assessed using the Shapiro–Wilk test. When assumptions of normality were met, comparisons between groups were performed using parametric tests (unpaired t-test or one-way ANOVA, as appropriate). Otherwise, non-parametric tests were used (Mann-Whitney test). Reproducibility was confirmed by independent experiments.

For IGFBP-1 and PRL secretion and NTA analysis of exosomes and microvesicles, data was analyzed using a two-way ANOVA to assess the effects of cell type and timepoint, as well as treated and non-treated. When significant effects were detected, pairwise comparisons were performed using Bonferroni’s post hoc test to correct multiple comparisons.

## Acknowledgments

We thank all patients who generously donated tissue and embryos for this study.

## Funding

Include all funding sources, including grant numbers, complete funding agency names, and recipient’s initials. Each funding source should be listed in a separate paragraph such as:

Carlos III Institute of Health grant PI21/00235 (IM)

Carlos III Institute of Health grant PI24/00722 (IM)

Carlos III Institute of Health grant PI21/00528 (FV)

Carlos III Institute of Health grant PI24/01784 (AMS, FV)

Spanish State Research Agency (AEI) grant CNS2022-135696 (FV)

Conselleria de Innovacion, Universidad y Ciencia CIAICO 2022/242 (FV)

European Commission Grant Agreement 101080219-2 (CS, FV)

Torres Quevedo Grant PTQ2022-012376 (MPF)

Generalitat Valenciana CIGE 2023 (CIGE/2023/217) (AMS)

## Author contributions

Conceptualization: SZ, MPF, CS, FV

Samples: SM, LQ, FR, JPG

Methodology: SZ, MPF, JGF, JMA, AQ

Data Analysis: SZ, MPF, AMS, CS, FV

Writing—original draft: SZ, MPF, AMS, CS, FV

Writing—review & editing: SZ, MPF, AMS, XS, IM, NP, CS, FV

## Data and materials availability

All data needed to evaluate the conclusions in the paper are present in the paper and the Supplementary Materials.

## Supplementary Materials

Please, see Supplementary materials included in a separate document

