## Supplementary material for "Modeling human embryo adhesion using a microfluidic platform"

Sofia Zaragozano *et al.*

**This PDF file includes:**

Figs. S1 to S4
Tables S1 to S4

Supplementary figure 1

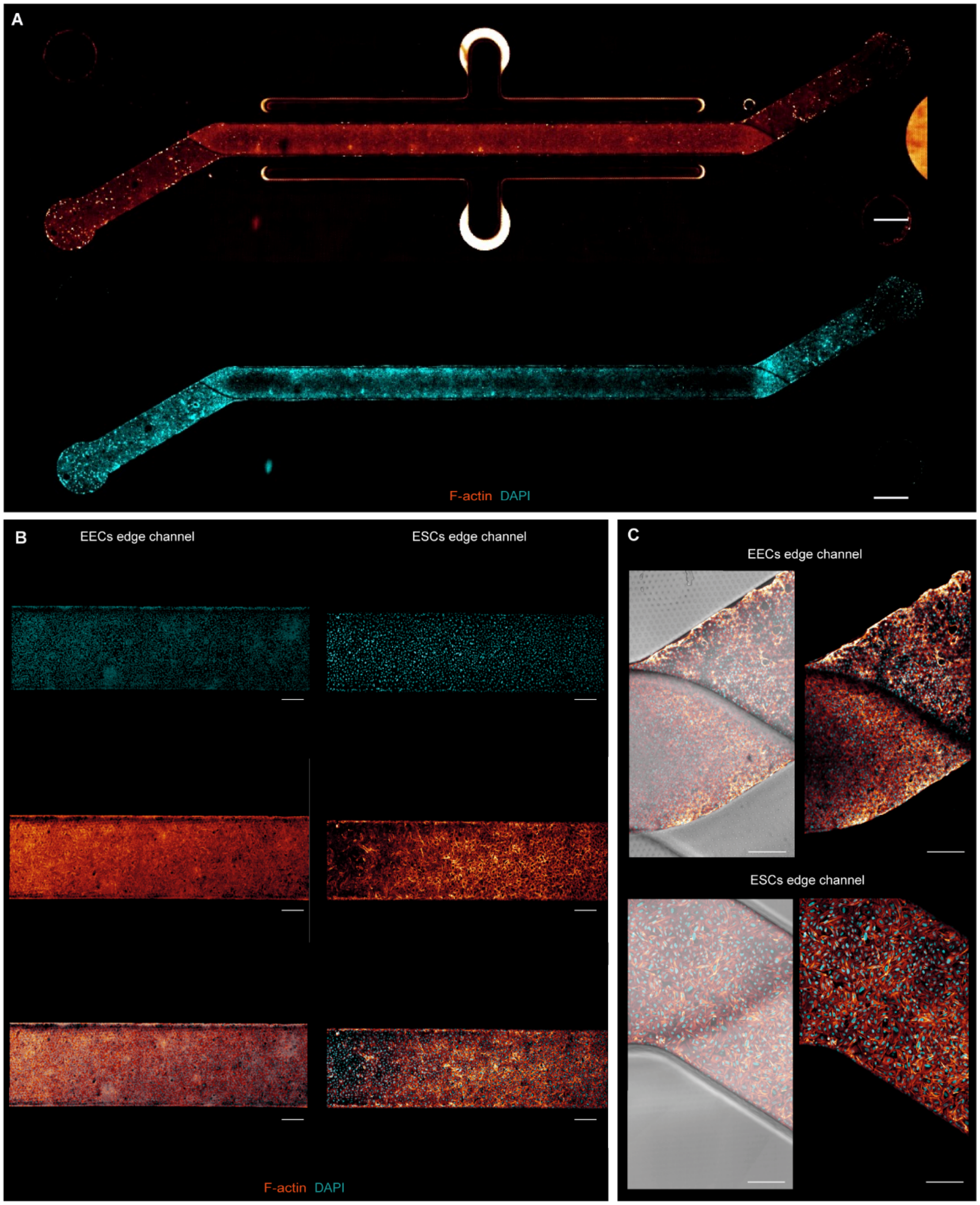


**Fig. S1. Morphological characterization across the microfluidic channels. (A)** Composite tile scan fluorescence image 8 days post-seeding, showing a fully confluent monolayer of organoid-derived endometrial epithelial cells and biopsy derived endometrial stromal cells. Scale bars, 1000 µm. **(B and C)** Higher magnification views of the center and edge channels of the chip. Cells nuclei are shown in blue, cytoskeleton (F-Actin) is shown in glow dark. Scale bars, 300 µm **(B)**, 200µm **(C)**.

Supplementary figure 2

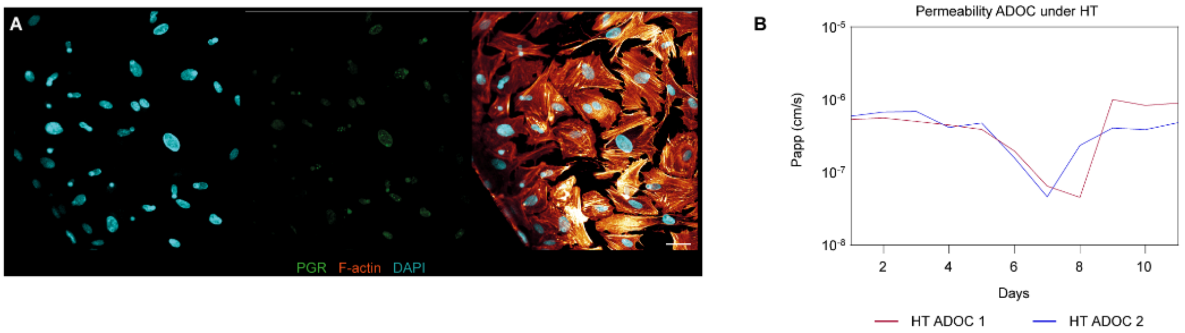


**Fig. S2. Hormonal response in ADOC. (A)** Representative immunofluorescence images demonstrating increased expression of progesterone receptors (PGR, green) in response to hormonal treatment, along with DAPI (blue) and F-actin (red) staining. Scale bars, 40 µm. **(B)** Changes in apparent permeability between the epithelium and stromal compartment of two different chips (n=2).

**Supplementary figure 3**

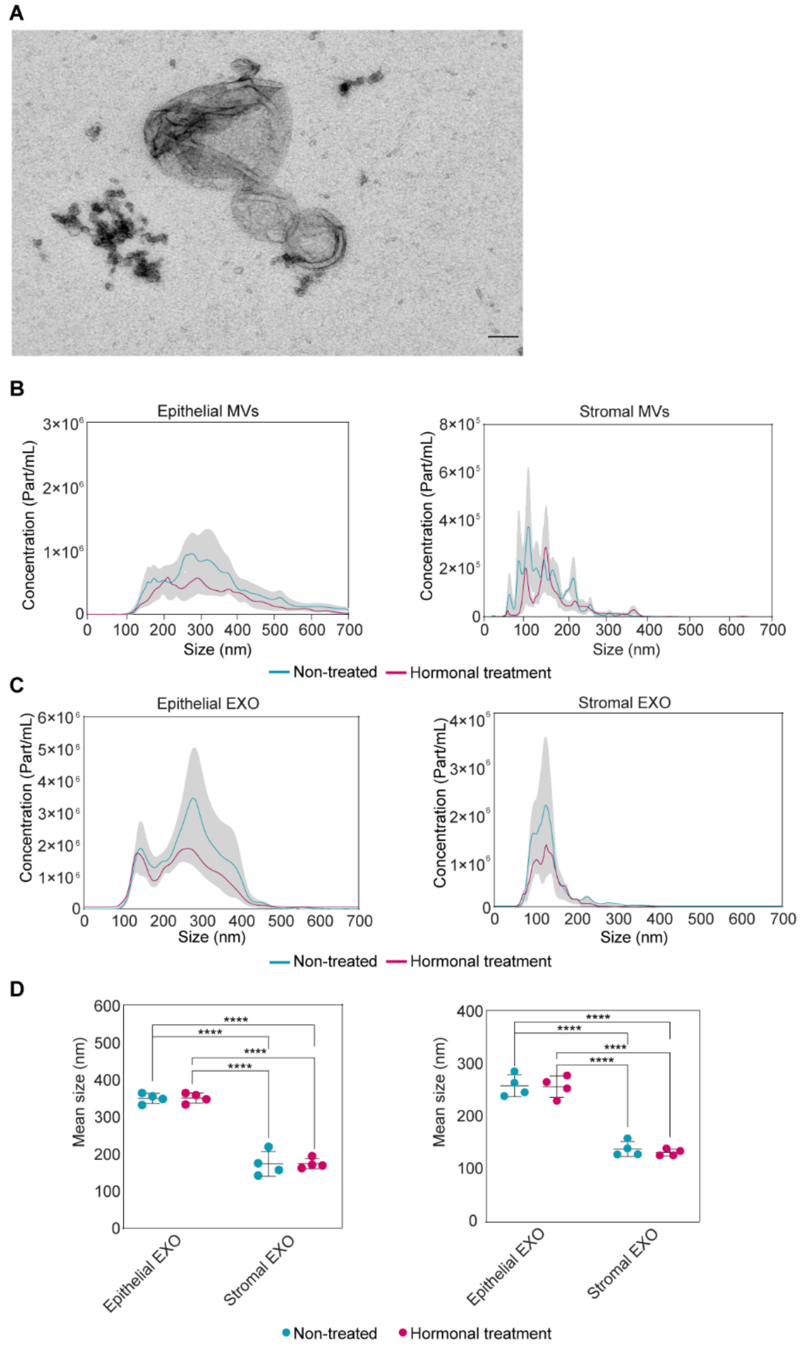


**Fig. S3. Morphology and nanoparticle tracking analysis of isolated EVs. (A)** Representative transmission electron microscopy (TEM) images showing apoptotic bodies (ABs) present in the medium of the chip. Scale bar, 100 nm. **(B)** Size distribution of microvesicles (MVs) as a function of particle concentration. **(C)** Size distribution of exosomes (EXOs) as a function of particle concentration. **(D)** Comparison of mean particle sizes for MVs and EXOs in media from hormone-treated and non-treated chips.

**Supplementary figure 4**

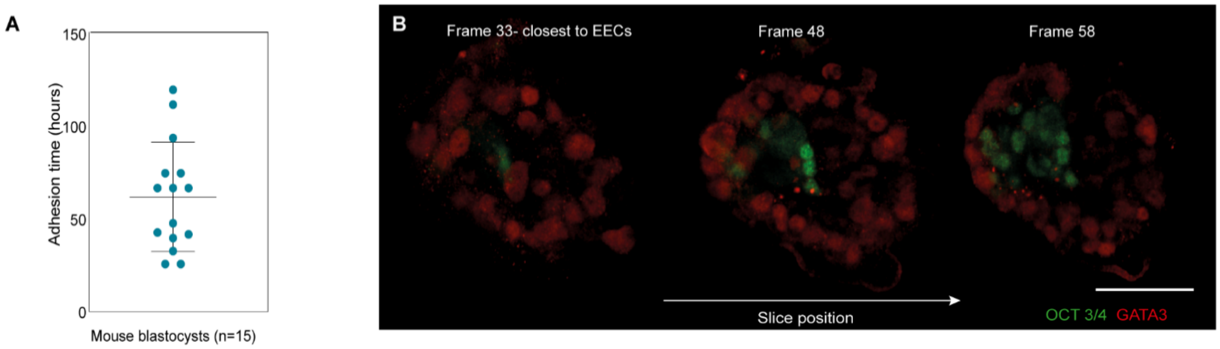


**Fig. S4. Adhesion timing and spatial orientation of mouse blastocysts in ADOC. (A)** Early adhesion time distribution of 15 embryos introduced into the chip, showing variability in the onset of adhesion. **(B)** Z-stack slices showing adhesion initiation by the trophectoderm. Left frame: plane closest to the epithelial layer (EECs). Right frame: plane farthest from the epithelial layer. Staining: ICM (OCT3/4, green) and trophectoderm (GATA3, red). Scale bar, 70µm

**Table S1. Endometrial epithelial organoid (EEO) medium composition**

| **Product** | **Concentration** | **Supplier** | **Catalog #** |
|---|---|---|---|
| DMEM/F12 | | Gibco | 11330032 |
| R-Sponding | 200 ng/mL | R&D systems | 120-38 |
| Noggin | 100 ng/mL | R&D systems | 6057-NG-100 |
| B-27 Supplement | 2% | Fisher scientific | 12587010 |
| N2 Supplement | 1% | Fisher scientific | 17502048 |
| Insulin transferrin selenium | 1% | Life Technologies | 41400045 |
| Penicilin /Streptomycin | 1% | Fisher scientific | 15140122 |
| Nicotinamide | 5 mM | Merck | 72340-100G |
| A83-01 | 0.5 µM | Merck | SML0788-5MG |
| N-Acetyl L-cysteine | 1.25 mM | Merck | A7250-50G |
| Epidermal growth factor (EGF) | 50 ng/mL | R&D systems | 236-EG-01M |
| Basic Fibroblast Growth Factor (bFGF) | 2 ng/mL | Fisher scientific | PHG0264 |
| FGF-10 | 50 ng/mL | Prepotech | 100-26 |
| p38 inhibitor | 10 µM | Merck | S7067 |
| Y-27632 | 10 µM | Merck | SCM075 |

**Table S2. Embryo medium composition**

| Product | Concentration | Supplier | Catalog # |
|---|---|---|---|
| DMEM/F12 | | Gibco | 21041025 |
| FBSi | 10% | Biowest | S181B |
| B-27 Supplement | 2X | Fisher scientific | 12587010 |
| N2 Supplement | 1X | Fisher scientific | 17502048 |
| Insulin transferrin selenium | 1X | Life Technologies | 41400045 |
| Penicilin /Streptomycin | 1% | Fisher scientific | 15140122 |
| N-Acetyl L-cysteine | 1.25 mM | Merck | A7250-50G |
| Essential Amino Acids | 1X | Gibco | 11130051 |
| No essential Amino Acids | 1X | Gibco | 11140050 |
| 17β-estradiol (E2) | 10 mM | Merck | E2758-250MG |
| Cyclic adenosine monophosphate (cAMP) | 0.5 mM | Merck | B7880-10MG |
| Medroxyprogesterone acetate (MPA) | 1 µM | Merck | M1629 |

From Rawlings, et al. *eLife* **10**, e69603 (2021). Reference (*15*) in main manuscript.

**Table S3. Primary antibodies**

| Antibody | Supplier | Catalog # |
|---|---|---|
| ZO-1 | Invitrogen | 33-9100 |
| Vimentin | Biotechne | AF2105 |
| Acetylated α-tubulin | Merck | T47451-25UL |
| PAEP | Abcam | ab270525 |
| Pan-Cytokeratin | Abcam | ab217916 |
| IGFBP-1 | Invitrogen | PA5-61-388 |
| Prolactin | OriGene | CF500719 |
| GATA3 | Invitrogen | 14-9966-82 |
| GATA4 | Invitrogen | 14-9980-2 |
| OCT3/4 | Santa Cruz | SC5279 |
| GATA 3 | R&D systems | AF2605 |
| PGR | Cell signalling technology | D8Q2J |
| hCG | Abcam | ab9582 |
| NR2F2 | Abcam | ab211776 |
| Pan-Cytokeratin | Abcam | ab217916 |

**Table S4. Secondary antibodies**

| Antibody | Supplier | Catalog # |
|---|---|---|
| Anti-mouse 488 | Invitrogen | A21202 |
| Anti-rabbit 555 | Invitrogen | A31572 |
| Anti-goat 647 | Invitrogen | AF1924 |
| Anti-rat 488 | Invitrogen | A21208 |
| Anti-rat 647 | Invitrogen | A48272 |
| Anti-mouse 647 | Invitrogen | A32787 |
| Anti-mouse 568 | Invitrogen | A10037 |
